# A human cytokine response atlas to reconstruct underlying gene regulatory networks

**DOI:** 10.64898/2026.07.27.740961

**Authors:** Patrick O’Connell

## Abstract

Circulating cytokines encode immune state, yet their pleiotropy and cell-type specificity make constructing a unified atlas of immune cell responses to them challenging. Here, I transformed a single-cell atlas of approximately 10 million human peripheral blood mononuclear cells from 12 donors exposed to 90 cytokines into a multiscale model of cytokine response. A GPU-accelerated implementation of dimension-scalable single-cell perturbation integration network (D-SPIN) allowed for the creation of a signed, directed model of 9.6 million cells, 1,634 immune regulatory genes, and 40 cellular programs. The gene networks and cellular programs span canonical cytokine pathways and lineage relationships and delineated cytokine-specific activation and repression across immune states. Beyond established circuitry, the model nominated candidate regulatory interactions and identified the mitochondrial antioxidant *SOD2* as a prominent hub of innate immune cell signaling. Next, I developed CytoCarto, a web application that projects cytokine profiles onto these networks to prioritize dysregulated programs, candidate effector genes, cellular contexts, and disease-associated signatures. In a proof-of-concept analysis I input the cytokine profile of a patient with mitochondrial encephalopathy, lactic acidosis, and stroke-like episodes (MELAS) undergoing an episode of sterile inflammation and found CytoCarto prioritized metabolically reprogrammed monocytes and *SOD2*, consistent with a role for mitochondrial redox signaling in innate immunity.

## Introduction

A longstanding question in immunology is how to reconcile the incompletely resolved and pleiotropic effects various cytokines have on immune cells. The study of cellular responses to cytokines is currently restricted to analyses of typically one or two cytokines at most due to feasibility and scaling issues when we consider there are >130 recognized human cytokines and chemokines as of 2016^1^. Efforts to address this have recently been undertaken and include an *in vivo* murine cytokine atlas^2^ and an *in vitro* human cytokine atlas^3^; both of which employ scRNA-sequencing of immune cells as a readout to capture cellular responses. While these large scale functional genomic projects have already advanced our knowledge of cellular cytokine responses, they have not resulted in the ability to construct a probabilistic model linking a particular cytokine milieu to a disease state, biological process, or responsible cell type.

Here, I reasoned that the rich, single-cell functional genomics data of 10 million human immune cell responses to 90 cytokines (generated by Oesinghaus et al.^3^) could be used to construct a gene regulatory network (GRN) which could in turn be refined into a probabilistic model to predict underlying biological responses in a patient with a particular cytokine signature in their plasma. In addition, such a model allows for the identification of novel cytokine signaling hubs and the construction of networks linking a cytokine(s) first to intracellular effectors (transcription factors, kinases, phosphatases, etc.) and then functional immune response programs. These networks can both recapitulate known signaling pathways and reveal novel genes and pathways underling cellular cytokine responses. To generate such a GRN from this dataset I made use of dimension-scalable single-cell perturbation integration network (D-SPIN)^4^ which takes as input a large functional single-cell genomics dataset and outputs a highly accurate GRN exceeding other computational approaches.

Clinical grade cytokine panels are more frequently being ordered for patients with undifferentiated inflammation, however, these are not easily interpretable even by expert immunologists, leading to, at best, an overly broad characterization of the flavor of inflammation present, and at worst, no characterization at all. A recent study has sought to address this through a large clinical database, but still was only able to classify broad immune dysregulation endotypes^5^. To address this limitation I reasoned the modeled response of a patient’s clinical cytokine profile could prove especially valuable in identifying the specific type of inflammation present whether it be an undiagnosed inborn error of immunity (IEI), inborn error of metabolism (IEM) with an immune dysregulatory component, alternative inflammatory state such as hemophagocytic lymphohistiocytosis (HLH), or just a more fine-grained understanding of the present inflammation.

## Materials and Methods

### Reference single-cell dataset and quality-controlled preprocessing

The Parse Biosciences human PBMC cytokine-stimulation atlas, provided as a raw-count AnnData (.h5ad) object with donor identifier, cytokine stimulus, and cell-type annotation was used as input to D-SPIN^3^. This dataset was cleaned, pre-processed, and annotated previously and these annotations were used for downstream analysis^3^. The model was fit globally rather than separately by cell type. Before feature selection and model fitting, cells annotated as plasmacytoid dendritic cells (pDC), innate lymphoid cells (ILC), hematopoietic stem/progenitor cells (HSPC), or plasmablasts were removed as the numbers were too few to satisfy suggested D-SPIN criteria of > 25 cells per condition.

For each retained cell *c* and gene *g*, raw count *x_cg_* was library-size normalized to 10,000 counts per cell and transformed as 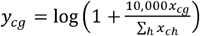. Here *h* indexes all genes measured in cell *c*. 2,000 highly variable genes (HVG’s) were then selected globally using scanpy (Supplemental table 1). These HVG’s were supplied to program-level model fitting. PBS-treated cells were labeled as controls, and the D-SPIN sample identifier was constructed as donor|cytokine. The original program-level model used 40 programs, 10 consensus orthogonal non-negative matrix factorization (oNMF) repeats, oNMF subsamples of 10,000 and 50,000 cells, square-root balancing, a maximum sampling rate of 2, k-means summarization, and 500 oNMF epochs^4^. The cell-type annotation was used to balance oNMF subsampling.

### GPU-optimization of D-SPIN

D-SPIN was extended with a CUDA/CuPy backend for the computationally intensive optimization steps. GPU acceleration was applied to both program- and gene-level pseudo-likelihood network inference, including estimation of the interaction matrix J and sample-specific response fields H, and to gene-to-program regulator discovery. Cell-level state matrices were processed in adaptively sized chunks to limit GPU memory usage, while model parameters and gradient calculations remained on the GPU. The CuPy implementation preserved D-SPIN’s sample weighting, regularization terms, Adam optimization, gradient monitoring, and backtracking behavior, and used double-precision arithmetic by default to improve agreement with the original implementation. The MATLAB and python-CPU backends were retained as alternatives, and upstream normalization, gene-program discovery by oNMF, discretization, and downstream output generation were unchanged. GPU-D-SPIN was used for the analysis presented herein.

### Program discovery and program-level D-SPIN network inference

Program discovery was performed by D-SPIN consensus oNMF on the normalized expression matrix restricted to the 2,000 HVG’s. Consensus discovery across 10 runs was used to stabilize the program basis. Program labels were assigned after fitting from the high-loading genes, local GO Biological Process/Reactome enrichment, and manual review. For each donor|cytokine sample, D-SPIN inferred a signed directed interaction matrix J^(^P^)^ among the 40 program states using pseudo-likelihood network inference with controls and batch information. The edge orientation was fixed as 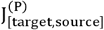. The program-network edge filter threshold was 0.02 during model construction. D-SPIN relative responses to PBS control were then calculated for every program and stimulus. Network modules were detected by Leiden clustering on filtered edges at resolution 1.0; a 0.05 network threshold was used for module discovery.

### Gene-level D-SPIN network and program-regulator determination

The gene-level model was fit with an initial list of 1,634 regulatory genes focused on transcription factors (TF’s), kinases, phosphatases, epigenetic regulators, translation regulators, proteostasis regulators, signal transductors, and more (Supplemental table 2). Genes were retained only if their nonzero fraction was at least 0.04, they were expressed in < 95% of all cells, and their variance exceeded 1e^-12^ after normalization. This prefilter prevents numerical instability in the D-SPIN heavy-tailed discretization, which clipped values at the 96^th^ percentile and also remove housekeeping genes with near universal expression. The gene model used directed pseudo-likelihood D-SPIN with step size 0.05 and L1 interaction penalty lambda = 0.01. It produced the signed directed gene network J^(G)^ and a gene-relative-response matrix.

To link the gene and program networks, gene and program state matrices were aligned by exact cell identifier. D-SPIN program-regulator discovery then estimated a gene-by-program matrix *B*, where 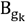 quantifies the signed association between program *k* and gene *g* in the fitted reference. Regulator fitting used 500 epochs, step size 0.02, and L1 interaction penalty 0.01. No threshold based solely on a patient’s observed cytokines was applied when generating *B*; it is a fixed reference object.

### Construction of gene and program network figures

The program-module network was generated from the 40 × 40 D-SPIN-inferred program interaction matrix. Interactions with absolute weight below 0.05 were removed before Leiden community detection (resolution = 1.0; random seed = 0), identifying five program modules that were subsequently assigned biological annotations based on their constituent programs. For visualization, programs were represented as module-colored nodes and interactions with absolute weight ≥0.15 were retained. Edge width and opacity were scaled by interaction magnitude. Node positions were determined using a module-constrained force-directed optimization balancing within-module cohesion, interaction-weighted attraction, and node repulsion.

The cytokine–program network was constructed from D-SPIN program responses relative to PBS. Responses were averaged arithmetically across 12 donors for each of 90 cytokines and 40 programs, and 479 cytokine– program connections with an absolute mean response ≥0.15 were displayed. Cytokines were grouped into 14 families exactly as in the original publication^3^. Edge width and opacity are proportional to the absolute donor-mean response.

The gene-module network was constructed from the D-SPIN gene-level interaction matrix and an exact-symbol annotation table containing gene-module assignments and functional classifications. For each gene pair, the finite, nonzero interaction from the nonredundant lower triangle of the matrix was retained, ranked by absolute interaction strength |J_ij_|, and the 2,500 strongest interactions were selected for plotting; genes with < 2 interactions were excluded. Edges were plotted if the absolute value of their weight was > 0.06. Node color represented the annotated gene-function class, while node area scaled with the square root of the number of displayed connections, 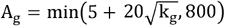, where k_g_ is the node degree. Line width and opacity increased with |J_iJ_|. Positions were calculated using a deterministic, interaction-weighted NetworkX spring layout with fixed module anchors to preserve separation among the five annotated modules.

The innate immune signaling network was generated by first restricting the annotation table and interaction matrix to genes assigned to the innate inflammatory signaling module. All within-module interactions were retained, and genes with < 1 edge were removed. Edges were plotted if the absolute value of their weight was > 0.05. Node colors used the same functional annotations as the complete network, all displayed genes were labeled, and node area followed 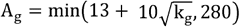. Positions were initialized circularly and optimized using an absolute-interaction-weighted spring layout, followed by radial compression, r_′_= tanh(r/r_80_) where r_80_ is the 80^th^ percentile of node distances from the network center. Edge sign, color, width, and opacity were encoded identically to the complete gene-module network.

The cytokine–gene–program effector networks were generated in Python from D-SPIN relative-response, gene-network, and gene-to-program regulator tables. For each selected cytokine gene and program responses were averaged across matching donor samples; target programs were specified directly. Up to 30 genes were initially ranked by their maximum absolute cytokine response, with genes classified as shared effectors when their second-largest response was at least 50% of the largest. Cytokine-to-gene edges and gene-to-gene edges were each individually thresholded for each network to prioritize readability. In multi-program figures, additional mediator genes were ranked by M_cgp_ = |R_cg_G_gp_|, where *R_cg_* is the cytokine response of gene *g* and *G_gp_* is its regulator score for program *p*; mediators were retained only when the sign of R_cg_G_gp_ agreed with the observed program response. Genes with fewer than two displayed connections were removed. Cytokines, genes, and programs were shown as diamonds, circles, and hexagons, respectively, and positioned using a deterministic interaction-weighted spring layout. Edge width and opacity were proportional to absolute effect strength. No direct cytokine-to-program edges were drawn.

### scRNA-seq analysis and UMAP density plots

The pre-processed Parse Biosciences scRNA-seq data did not contain UMAP embeddings and was also required for calculating program responses on a per donor, cell type, and cytokine family basis. Due to these factors a more heavily processed scRNA-seq dataset was required and I perform standard analysis of using the Seurat pipeline^6^ with the BPCells approach considering the massive size of the dataset. Data processing and analysis was performed on the Mount Sinai Minerva HPC. A sketch assay was created using 200,000 cells and full analysis of the data including UMAP calculation was performed on this sketch object. The sketch data was projected back onto the entire 10 million cell dataset and downstream analysis including UMAP figure generation were performed with the entire dataset.

Per-cell D-SPIN program scores were stored as Seurat metadata, while genes assigned to each annotated gene-network module were grouped into module gene sets and summarized into one activity score per module and cell. These program and module scores were then treated as continuous cell-level features and aligned to the corresponding cells in the fixed UMAP embedding. For each feature, the R package Nebulosa was used to calculate a two-dimensional score-weighted kernel density over the UMAP coordinates. The resulting density was mapped back to each cell and displayed using rasterized points. Consequently, color indicates UMAP neighborhoods enriched for high program or module activity rather than raw cell abundance or expression in an individual cell.

### Design of circos plot to visualize program responses across cell types and cytokine stimulations

The circos plot was constructed in R using the Circlize package. For each donor *d*, cell type *c*, cytokine *k*, and D-SPIN program *p*, the cell-level scores were averaged as 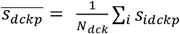,and the matched PBS response was subtracted: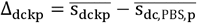. The plotted effect was the equally weighted donor mean,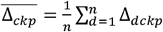 .A two-sided one-sample paired t-test evaluated whether donor - level deltas differed from zero, 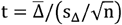, with *n*−1 degrees of freedom, and *p* values were Benjamini–Hochberg corrected within each cell type. Cell types formed the circular sectors, cytokines were ordered within each sector by cytokine family, and radial lanes represented programs P0–P39. Only effects with *q*_BH_ < 0.05 were shown; dot color encoded the signed delta, while dot size scaled with capped -log_l0_(FDR). For each cytokine responses were averaged to obtain a 40-program delta vector for every cell type, and similarity between cell types was calculated using Pearson correlation. Inner links were retained only for *r* > 0.95 or *r* < −0.3, with link color indicating correlation sign and line width and opacity increasing with |r|.

### Construction of the CytoCarto application

CytoCarto was implemented as a Python workflow comprising three separate processing scripts. The workflow accepts normalized plasma cytokine measurements with optional patient age, sex, race, and ethnicity, projects the observed cytokine state onto a fixed D-SPIN peripheral-blood program and gene network, identifies enriched cellular contexts using the 10-million-cell PBMC reference, and generates human disease, mimic, and perturbation evidence. Outputs are designed to be hypothesis-generating and are not diagnostic.

### Cytokine normalization and covariate-aware response aggregation

Input cytokine names were first harmonized to the D-SPIN immune map. For each cytokine *i*, the reported concentration *x_i_* was normalized to the upper laboratory reference limit *r_i_*. Censored values below the limit of detection retained their original qualifier for display while using the established censored-value handling for scoring. Only elevations contributed to the network seed:

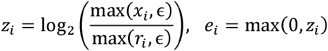

where *z_i_* is the log2 fold elevation, *e_i_* is the non-negative cytokine-elevation score, and ∈ = 10^−9^ prevents division by zero. Soluble IL-2 receptor alpha was retained as a clinical analyte but excluded from D-SPIN network seeding because it is not a cytokine node in the D-SPIN network.

The reference atlas contained donor-level cytokine stimulation responses allowing CytoCarto to take into account the age, sex, race, and ethnicity of the query subject (patent). Donor (subject whose cells were sequenced as part of the human immune cytokine atlas) observations were weighted by similarity to the patient’s available demographic variables. Age similarity was weighted using an exponential kernel with a 30-year scale; sex mismatch received weight 0.6; race or ethnicity mismatch received weight 0.7; and missing donor demographic information, when the corresponding patient variable was supplied, received weight 0.85. If age was available for the patient but unavailable for a donor, the age component was set to 0.8. The final donor weights were normalized to sum to one before computing weighted mean gene-response vectors for each assayed cytokine.

### D-SPIN program scoring

For each cytokine-associated program response vector 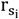, the patient program seed was calculated as the elevation-weighted mean response across the observed cytokines:

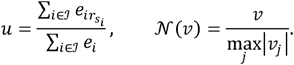

Here, *J* is the set of recognized elevated analytes, *u* is the initial 40-program patient state, and *N* rescales a nonzero vector to unit maximum absolute magnitude. The signed program state was retained for subsequent network and public-evidence analyses; the reported program-elevation display score was *N*(max(u,0)).

The D-SPIN program interaction matrix J^(P)^ used target-by-source orientation, such that 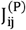 denotes the effect of source program *j* on target program *i*. Rows were normalized by the sum of absolute off-diagonal weights, preserving edge direction and sign:

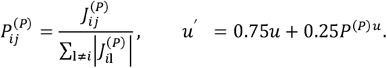

The final signed program score was *N*(*u*^*′*^). Thus, direct cytokine-derived program activity retained 75% weight, while one directed network-propagation step contributed 25%. Positive and negative network edges therefore represented activating and opposing relationships rather than unsigned connection strength.

### Gene-level scoring and propagation

Direct cytokine-to-gene associations were assembled into a gene seed vector *d*, where each gene was assigned the strongest matched cytokine elevation among its mapped inputs. Program-derived gene regulation was calculated using the program-to-gene loading matrix *B*:

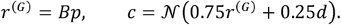

where *p* is the final signed program vector, *r*^(G)^ is the program-derived gene-regulatory state, and *c* is the combined signed gene seed. The 75:25 weighting prioritized program-mediated inference while preserving direct evidence from elevated measured cytokines. The gene network was then propagated once using the signed, directed D-SPIN gene interaction matrix:

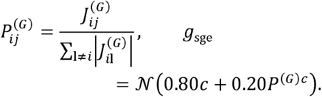

Here, 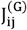 is the directed influence of source gene *j* on target gene *i*. The signed gene vector was used for direction-sensitive evidence retrieval. A non-negative display vector was generated separately by clipping negative values after propagation; clipping was not applied to disease or perturbation matching.

### Exact cell type dysregulation enrichment

Cellular context was determined by comparing the patient’s signed 40-program vector with every cell in the 10-million-cell PBMC reference program representation. For cell *c*, with program vector a_*c*_, cosine similarity to the patient vector *p* was calculated as:

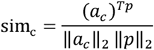

The highest-scoring 1% of cells were designated the dysregulated-cell candidate set. Exact row-order and cell-identifier agreement were required between the per-cell program matrix, the program observation names, and cell metadata.

For each cell type *t*, enrichment was calculated from the fraction of all reference cells belonging to that type and the fraction of top-matching cells belonging to that type:

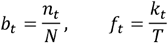

Where *b_t_* is the background frequency of each cell type in the full 10 million cell atlas, *n_t_* is the number of cells in that cell type, *N* is the total number of cells in the atlas,*f*_*t*_ is frequency of a cell type among cells most similar to the query profile, *k*_*t*_ is the number of top matching cells for each cell type, and *T* is the total number of retained top matching cells. The cell type fold enrichment is then calculated as: 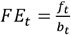.

### Human disease, mimic, and perturbation evidence layer

The disease/mimic layer is a human-only retrieval and evidence-ranking module. This layer asks: *does the patient’s inferred signed gene state resemble the genes linked to known human diseases, mimics, pathways, regulatory programs, or cell contexts?* It consumes the already computed signed gene state, signed program state, D-SPIN gene network, gene-to-program matrix, and cell-type table; it does not modify cytokine, program, gene, or cell-type scores. It uses the ChEA-KG (a signed transcription-factor network) database to perform directed-alignment of this large GRN with the subject’s cytokine-profile-derived query GRN. Records labeled mouse, mixed-species, or unknown species are excluded. Patient signatures are not transmitted during cache-only scoring. Enrichr-KG is treated as a heterogeneous term-gene knowledge graph, not as a gene-regulatory network.

Multiple databases were included and are listed here with their purpose. Direct disease evidence is obtained from human DisGeNET, Jensen DISEASES, and GWAS Catalog terms. Human Phenotype Ontology and pathway resources (GO Biological Process, Reactome, KEGG Human, and WikiPathways Human) provide supporting interpretation. ChEA 2022, TRRUST, and ARCHS4 transcription-factor co-expression provide regulatory support; Tabula Sapiens, HuBMAP ASCT+B, Descartes, and Human Gene Atlas provide cell-context support. LINCS CRISPR knockout signatures provide directional perturbation support. STRING protein-protein interactions and a symmetrized ARCHS4 co-expression projection are used only as unsigned proximity graphs. CCLE, Achilles, and acquired-disease terms provide mimic-context evidence. Human Perturb-Seqr records are restricted to explicit gene perturbations and are reported as mechanistic phenocopy or reversal evidence.

For an unsigned term *T*, direct enrichment uses absolute gene magnitude,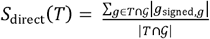. For directional signatures, values are multiplied by the signature direction before averaging. The top 200 direct-scoring terms per library are assessed with 2,000 degree- and term-size-matched permutations. Degree strata are calculated from the combined incident connectivity of D-SPIN, ChEA-KG, STRING, and ARCHS4 graphs. An empirical P value is calculated as (1 + number of null scores greater than or equal to the observed score)/(1 + number of permutations); adaptive additional permutations are used for direct disease and cancer-context terms when needed. Benjamini-Hochberg correction is performed within each library. Duplicate signatures from the same study do not multiply a disease’s evidence because the best empirical P value per resource is retained before combining sources.

Signed network support is calculated independently at propagation depths 0 through 3 through both D-SPIN and ChEA-KG networks after diagonal removal and absolute-row normalization. The unsigned multi-view retrieval embedding is built from absolute D-SPIN and ChEA connectivity, STRING PPI, and ARCHS4 co-expression; a separate four-channel signed sensitivity embedding preserves positive/negative incoming/outgoing connectivity. These network, manifold, PPI, and co-expression measures are reported separately as support and candidate-module retrieval measures. They are not included in the primary named-disease score unless a leave-one out validation has shown a positive bootstrap 95% confidence interval for improvement over direct enrichment.

Named disease terms are normalized to MONDO. The primary named-disease P value combines only independent direct disease-library empirical P values by equal-weight ACAT (Aggregated Cauchy Association Test):

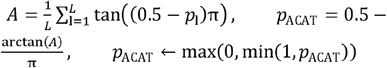

Where *L* is the number of distinct disease resources, *ℓ* is the index of these resources, and *p*_*ℓ*_ is the empirical disease-enrichment P value from resource *ℓ*. The evidence score is -*log*_l0_(*max* (*p*_*ACAT*_ 10^−300^)) and Benjamini-Hochberg correction is then applied across ranked named diseases. Hypotheses are classified from source/ontology names as monogenic/genetic disease, acquired inflammatory or autoimmune mimic, infection/sepsis-like mimic, cancer-associated mimic, or other disease evidence. Mimics are ranked separately from named genetic disease hypotheses.

## Results

### Construction of a program network of PBMC cytokine responses

D-SPIN works by constructing both a GRN and a program network from the input data and then identifying linkages between how the GRN influences various program responses. By incorporating a large number of perturbations (cytokine stimulations in this case) it is able to identify otherwise hidden gene interactions which would be masked in a GRN constructed using other methods^4^. I reasoned that by providing as input a 10 million cell scRNA-seq data set of human PBMCs subjected to stimulation with 90 different cytokines *in vitro*, a highly accurate model of human peripheral immune cell responses to cytokine stimulation could be constructed. I used the Oesinghaus et al., dataset which contains the above-mentioned data and encompasses 12 healthy donors of varying ages^3^. After removal of rare cell types, the top 2,000 highly variable genes (HVG’s) were calculated and this was used as input to construct the program level network. The number of programs was pre-set at 40 per D-SPIN program recommendations and for the sake of interpretability. The programs underwent semi-manual annotation based on genes identified as important in each (Fig. 1A). These programs compose specific immune cell subsets, general cellular processes, or a combination of both. The 40 programs were clustered into modules and resolved into five modules representing immune cell types and states (Fig. 1B). Programs within a module positively reinforce other programs of the same module while opposing programs from other modules (Fig. 1B). For example, multiple programs within the “lymphocyte homeostasis” module repress the most general “myeloid homeostasis” program “P0-Myeloid cells” (Fig.1B).

**Figure 1:**
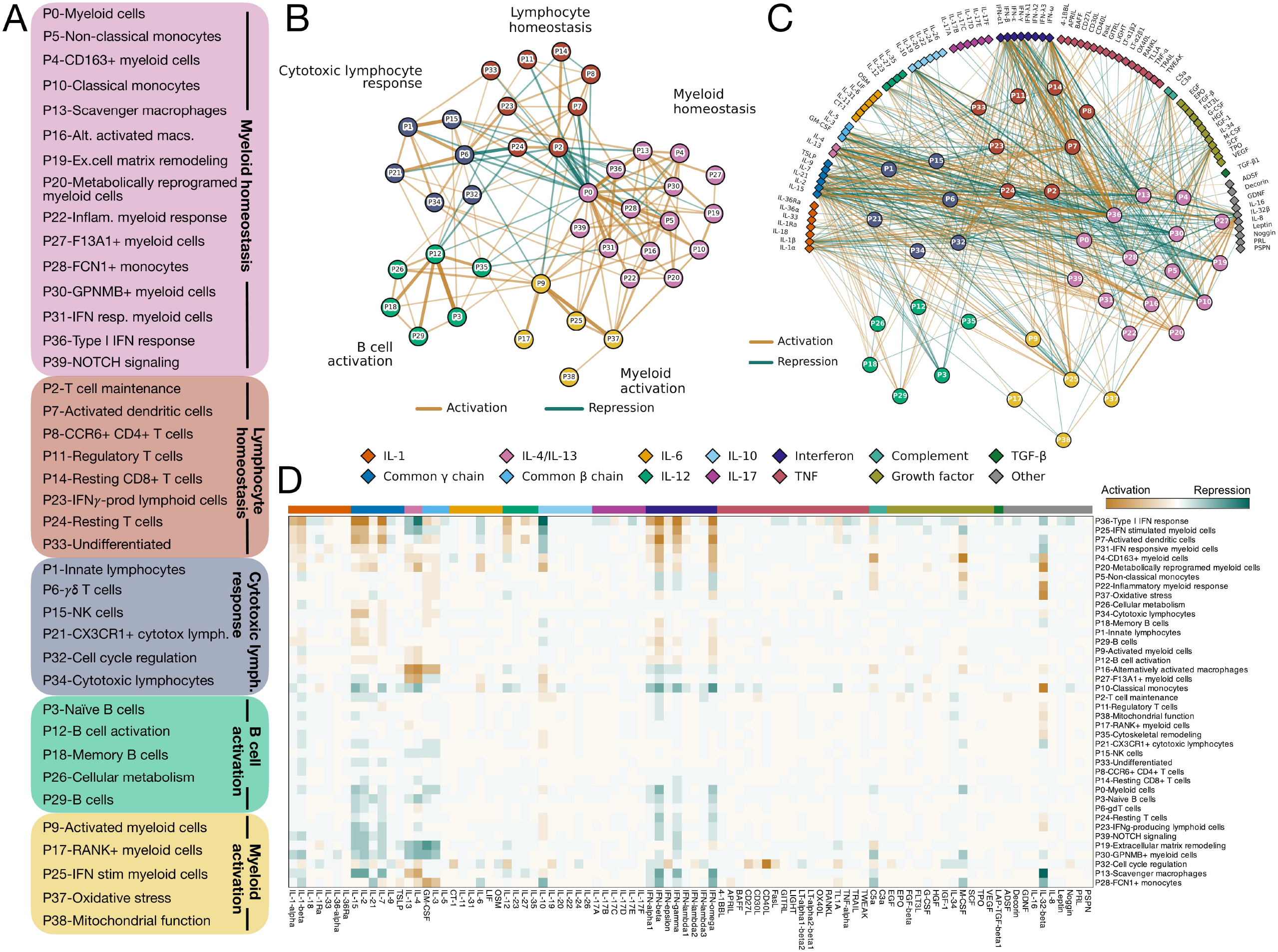
A program network of human PBMC cytokine responses. A. List of all 40 programs identified by oNMF and semi-manually curated. Programs are grouped by modules as determined by Leiden clustering. B. Network of all programs naturally grouped by modules. Brown edges indicate activation or positive reinforcement between programs while green edges represent repression. Edge width and opacity are scaled to the strength of the connection between nodes (programs). C. Program network including all cytokines (grouped and colored by family) and showing interaction each cytokine has on each program. Interactions (edges) are scaled in color and opacity similar to (B). Between node edges are not shown for clarity. D. Heatmap of program responses to each of the 90 cytokines with brown indicating activation of that program and green representing repression.

I next examined how each cytokine activates or represses these programs. Similar to Oesinghaus et al., I found that a select few cytokines (ie. IL-32β, IL-2, IL-15, IL-1β, type I and type II interferons, just to name a few) had pronounced effects, while many other cytokines had a more limited effect on each

### Mapping D-SPIN program responses to single cells

The program network D-SPIN constructs takes into account each cells type and then assigns a program response score for each of the 40 programs to every cell. Using these single-cell responses I next mapped program responses for each cytokine across all immune cell subsets. UMAP dimension reduction of the 10 million cell dataset shows well resolved immune cell subsets in a distribution one would expect for scRNA-seq data of human PBMC’s (Fig. 2A). Using this cellular atlas I then map program module scores back to single cells and visualize this with kernel density estimation. Examining a few programs, we find that P0-Myeloid cells align well with the CD14 Monocyte and CD16 Monocyte populations, myeloid cells in general are the only cell types program (Fig. 1C, D). Similarly, some programs were strongly influenced by cytokine stimulation (P36-Type I IFN response, P10-Classical monocytes, and P32-Cell cycle regulation) and others had limited responses (P17-RANK+ myeloid cells, P8-CCR6+ CD4+ T cells, and P39-NOTCH signaling) (Fig. 1C, D). The program network also shows, as expected, that many of the most potent cytokines simultaneously activate and inhibit opposing programs. For example, IL-32β activates an inflammatory myeloid response while inhibiting IFN responses and a scavenger macrophage program and IL-4 promotes an alternatively activated macrophage state while suppressing type I IFN responses (Fig. 1D). Critically, this same pattern of IL-32β was also observed by Oesinghaus et al. using a complementary approach^3^. Additionally, on a global level, myeloid cell programs show the most pronounced responses to various cytokine stimulations which is expected given these are innate immune cells (Fig. 1C, D).

**Figure 2:**
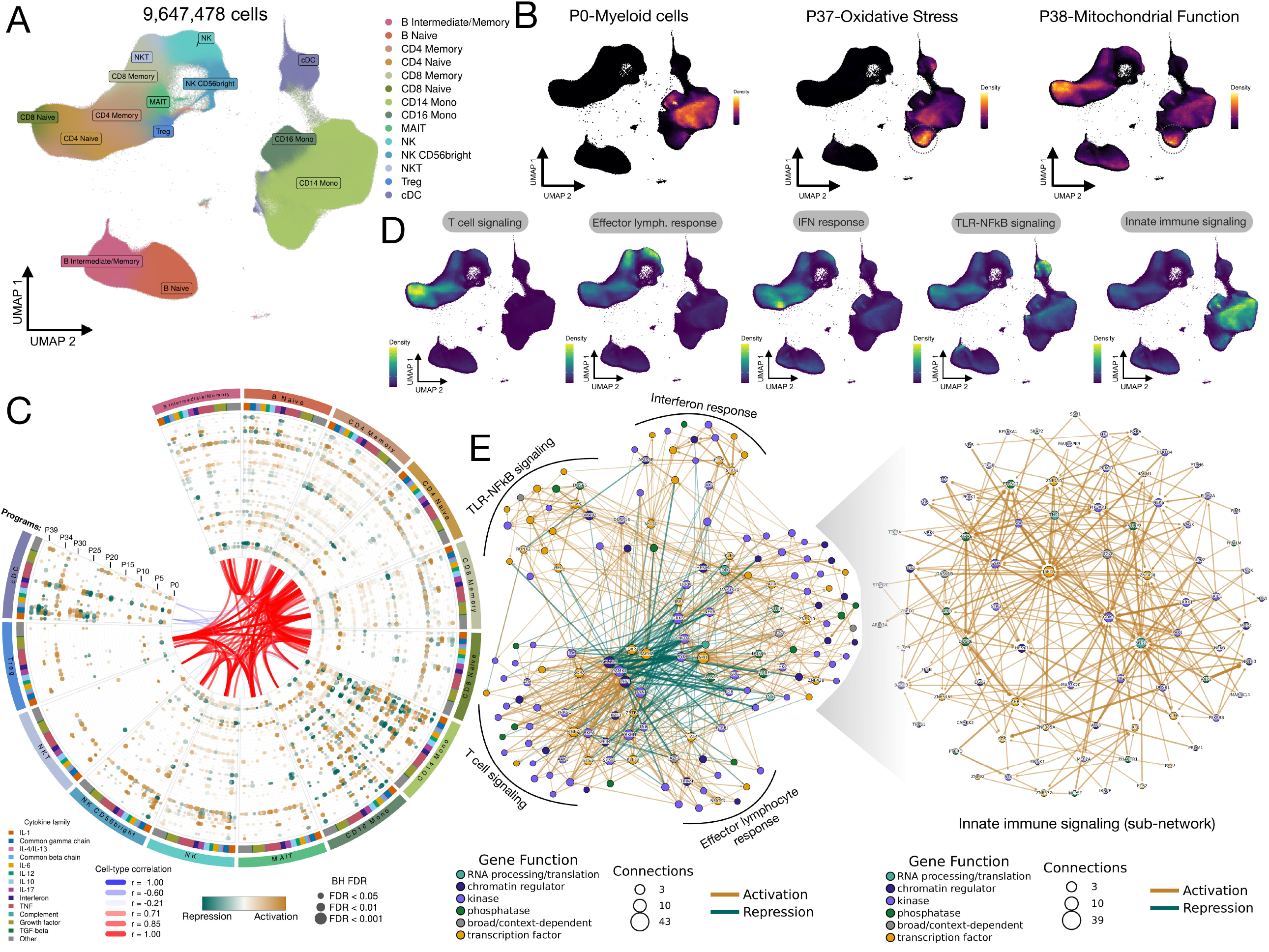
Program network and GRN responses mapped to immune cell subsets. A. UMAP of the full 10 million cell dataset used for D-SPIN analysis with cells colored by subset as determined by Oesinghaus et al^3^. B. Kernal density estimate of various program activities overlaid onto the cellular UMAP. C. Circos plot of program responses to each cytokine on an immune cell subset-specific basis. Cell types are indicated on the outermost ring with the next ring colored by cytokine family (and cytokines are ordered within that family the same as in Fig. 1D. The next ring contains a stacked dot plot with each dot a program response scaled in size by BH FDR and in color by strength of program response. Programs are arranged linearly from 0-39 from inside to outside. The innermost portion of the figure shows prominent negative and positive correlations between each cell type|cytokine pair. Positive correlations are in red and negative correlations are in blue with line scaled by width and opacity relative to correlation strength. D. Kernel density estimates of each gene module overlaid on the cellular UMAP. E. Left, subnetwork of the D-SPIN GRN showing the most highly connected regulatory genes. Nodes are scaled by number of connections and edges are scaled in width and opacity by the magnitude their strength. Nodes are colored by function. Right, subnetwork of the Innate immune signaling gene module showing only genes with >1 edge.

showing the P37-Oxidative Stress response, and the broad cellular function program P38-Mitochondrial Function is found in most cell types (Fig. 2B). Careful inspection reveals that a subset of CD14 monocytes (outlined) have particularly high scores for both the P37 and P38 programs, suggesting these are a particularly metabolically active subset of cells (Fig. 2B).

For a more quantitative and holistic view of program responses across each cytokine and cell type I used the single-cell level program responses to construct a cellular program response map (Fig. 2C). Here we see CD14 Mono, CD16 Mono, and cDC had the largest pan-cytokine responses amongst all cell types (Fig. 2C). NKT cells and Tregs had the most limited cytokine responses consistent with an orthogonal analysis of the same dataset^3^ (Fig. 2C). Some programs, in particular P32-Cell cycle regulation, had a fairly uniform response across all cell types and cytokines, fitting with an established role for cytokines as general regulators of immune cell proliferation^7^. When we look at how closely a given cytokine response correlates among cell types we note broad agreement among similar cell lineages with a few notable exceptions including IL-11 responses in cDCs and various lymphocyte subsets (Fig. 2C, Supplemental Fig.1 and Supplemental table 3). This is potentially explained by the ability of IL-11 to induce a *parfor* function in MATLAB. Jiang et al. use as many as 2 million cells in D-SPIN, but a dataset as large as 10 million cells has never been attempted (J. Jiang, personal communication). Even with parallelization to 64 during GRN generation could be sped up by running on a GPU. I took advantage of the python package CuPy^10^ to allow for streamlined porting to a GPU with limited changes t o the larger program architecture. This resulted in congruent results comparing the native D-SPIN python gene network construction to a GPU optimized version. A test run of 500,000 cells was performed to compare both approaches and run time on the program network calculation was equivalent (Supplemental Fig. 2A, B) with an approximate

### A cytokine-driven gene regulatory network of human PBMC’s

Using a curated list of genes known, or believed, to be critical to immune cell function, I leveraged D-SPIN to generate a gene regulatory network representing cytokine responses in human PBMC’s. Leiden clustering identified five distinct gene modules (T cell signaling, Effector lymphocyte response, IFN response, TLR-NFκB signaling, and Innate immune signaling). I took the genes composing each module and generated a module score proinflammatory IL-1β response in cDC’s^8^ while driving Th2 polarization in CD4^+^ T cells^9^.

### GPU optimization of D-SPIN

D-SPIN provides a function to take in a list of regulatory genes and output a highly accurate GRN with takes into account each perturbation to the system (cytokines in our case) to reveal previously masked gene interactions^4^. Given this task requires repetitive calculations over every cell in the dataset, run times become prohibitively long even with the included option of parallelization using the cores and 1.2 Tb of memory, D-SPIN could not complete the GRN calculation after running for multiple days, therefore, I reasoned the large matrix calculations D-SPIN performs 20-fold speed up on the gene network calculation (Supplemental Fig. 2B). This speed up is even more dramatic with larger numbers of cells where I found an 80-fold speed up when 2 million cells were run with the GPU pipeline versus the MATLAB pipeline (Supplemental Fig. 2C, D). The full 9.6 million cell Parse Biosciences dataset was able to run through gene-level D-SPIN with 1,634 regulatory genes in 8 hours on a NVIDIA H100-SXM5-80GB GPU.

which, when mapped back to the full 10 million cell UMAP, reveal expected distributions (Fig. 2D). While some modules mapped to specific cell subsets (T cell signaling and Effector lymphocyte signaling) other, more response-based modules (IFN response, TLR-NFκB signaling, and Innate immune signaling) broadly mapped to multiple cell types (Fig. 2D).

To explore this GRN further I constructed a network using the most connected genes (Fig. 2E). Genes in each of the five modules naturally grouped together, but considering this GRN reflects many cell types, a few genes are more central and cross-talk to many modules, for example, *REL, BACH2, FOXP1, FYN, SYK, SP1*, and the less studied *GRK3*^11^ (Fig. 2E). In general, nodes within a module positively reinforce each other and the strongest repressive interactions are between hub nodes of opposing cell types, such as T cell signaling and Innate immune signaling (Fig. 2E), similar to what was noted on the program-level network (Fig. 1B). It was reassuring to see D-SPIN correctly placed genes in the network as evidenced by, for example, the central placement of *ETV6* in the IFN response network (Fig. 2E). *ETV6* has been shown to control monocyte IFN responses^12^, but a complete mechanism explaining this remains elusive. This GRN shows strong positive interactions between *STAT1, ALPK1*, and *SP100* with *ETV6* which suggest possible mechanistic partners for further evaluation. I explored the Innate immune signaling module in more detail by generating a network representation of the most important nodes and edges in that subnetwork (Fig. 2E). This network is composed of only positive regulation between nodes which is likely secondary to the method of perturbation (cytokine stimulations) used to construct the GRN. An approach that more uniformly disrupts many innate immune cell functions, such as genome-wide Perturb-seq, would in theory address this. Regardless, this network reveals multiple expected findings such as *SP1* being the most critical regulator of innate immune signaling and co-regulation of *AHR* and *KLF6* by *PRDM1*^13,14^ (Fig. 2E).

Interestingly, *SOD2* emerged as one of the most critical hub genes after *SP1* (Fig. 2E). no specific IFN*γ* program was identified, and the strongest IFN*γ* response noted (Fig. 1D). the P36-Type I IFN response program contains Our human network closely mimics the related *SOD2* encodes manganese superoxide dismutase which plays an important role in controlling mitochondrial oxidative stress^15^. SOD2 has primarily been studied in the context of cancer and is not recognized as a master regulator of innate immune responses, however, there is a report showing *SOD2* plays a role in innate immune responses to bacterial infections in zebrafish via ROS regulation^16^. A recent theory has emerged that mitochondria not only play a critical role in cellular metabolism and energy production, but in fact function as signaling hubs coordinating a diverse array of cellular processes including immunity^17,18^. The centrality of *SOD2* in our comprehensive cytokine-response network supports this view.

### Reconstructing functional cytokine-program networks

One of the greatest strengths of D-SPIN is its ability to simultaneously link effectors (cytokines in our case) to genes to programs^4^. Such investigations have the potential to uncover both established and undiscovered signaling pathways connecting an effector(s) to cellular program(s) of interest^19^. I investigated a select few cytokine-program networks to assess if the results are consistent with prior analyses and to explore how a single cytokine and can exert opposing effects on different programs. I first investigated how IFN*γ* and IFN⍰induce the P36-Type I IFN response program (Fig. 3A). Jiang and Thomson investigated a similar network, albeit applying D-SPIN to an *in vivo* murine cytokine response atlas (T cells only) and an IFN response program^19^. Of note, in the human D-SPIN model

**Figure 3:**
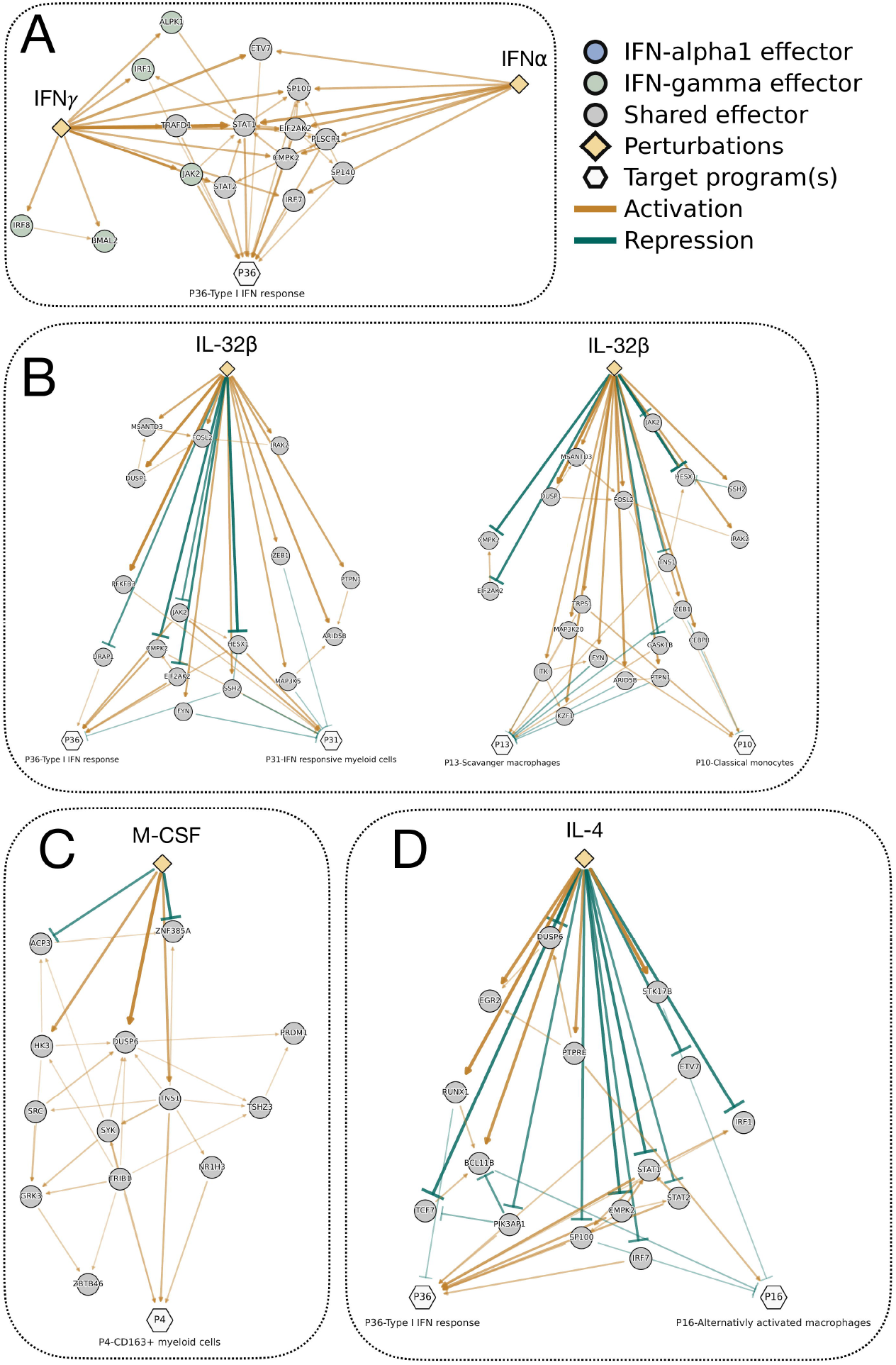
Reconstruction of GRN’s linking cytokines to immune programs. A. Cytokine-program network of IFN_γ_ and IFN⍰ and the P36-Type I IFN response program. Edges are scaled in width and opacity by magnitude of association. Node size is fixed and any node with < 2 edges is removed. B. Left, cytokine-program network for IL-32β and the P36-Type I IFN response and P31-IFN responsive myeloid cells. Right, cytokine-program network for IL-32β and the P13-Scavanger macrophages and P10-Classical monocytes programs. C. Network of M-CSF activation on the P4-CD163+ myeloid cells program. D. Network of IL-4 regulation on P36-Type I IFN response and P16-Alternativly activated macrophages programs.

murine network with IFN*γ* activating the program through *IRF1, JAK2, STAT1*, and *STAT2*, while IFN⍰activates the program through *STAT1* an d *IFR7* (Fig. 3A).

Next, I examined how IL-32β regulates various programs, considering this pleotropic and understudied cytokine was noted by Oesinghaus et al.^3^ to have particularly profound and divergent effects. IL-32β was noted to inhibit IFN responses by Oesinghaus et al. and also inhibited IFN-related programs in the D-SPIN model (Fig. 1D). Examining the (Fig. 3B). I also observed that IL-32 had opposite effects on the P13-Scavanger macrophages and P10-Classical monocytes programs (Fig. 1D). Exploring the network, we see it accomplishes this by inducing negative regulators of scavenger macrophages (CD206^+^, “M2-like”) such as *PTPN1*^20^, *IKZF1*^21^, and *FYN*^22^ and activating positive regulators of the classical monocyte state such as *FOSL2*^23^ and *TRPS1* (which suppresses GATA-1, a known suppressor of monocyte lineage commitment^24^) (Fig. 3B).

Since M-CSF is well-known to polarize macrophages to an “M2-like” state^25^ and induce CD163 expression, I evaluated the GRN between this cytokine-program dyad. It is notable that genes encoding the well-established pathways governing M-CSF macrophage polarization (*PI3K, AKT, RAS, ERK*, and others) are not included in this network (Fig. 3C). Instead, D-SPIN identified that M-CSF acted through *TRIB1, TNS1*, and *NR1H3* to drive the P4-CD163+ myeloid cells program (Fig. 3C). While *TRIB1* is a well-established positive regulator of “M2-like” macrophages^26^, *TNS1* has limited evidence linking it to M2 polarization^27^ and there is conflicting evidence for *NR1H3*.

Finally, I explored the opposing effects of IL-4 on the P36-Type I IFN and P16-Alternativly activated macrophages programs. IL-4 inhibits cytokine-effector network, it appears to accomplish this inhibiting direct IFN effectors (ie. *JAK2* and *EIF2AK2* [AKA PKR]) and activating possible inhibitors of the IFN response (ie. *FYN, MAP3K5*, and *ZEB1*), although the latter inhibitory relationships have not been well described and may represent novel signaling mechanisms

IFN signaling via repressing activators of the IFN response such as *STAT1, STAT2, IRF1*, and *IRF7* (Fig. 3D). Interestingly, it appears that IL-4 inhibition of *SP100* is critical for driving these diametrically opposed responses given *SP100* is a known interferon stimulated gene^28^ and was found to inhibit the P16 program (Fig. 3D). The later finding is not supported by prior research, but members of the SP protein family are recognized to control macrophages functions and *SP100* has been found in increased levels in “M1-like” macrophages^29^.

### Leveraging the D-SPIN cytokine regulatory network to assist in the interpretation of clinic cytokine panels

Physicians are more frequently ordering clinical-grade cytokine panels to better classify the flavor of a patient’s inflammation, to attempt to differentiate autoinflammation from true infection, and to inform therapeutic options. However, these cytokine panels remain challenging to interpret even for experienced clinical immunologist and rheumatologists^5^. I reasoned that the rich GRN and program regulatory networks generated by D-SPIN on this cytokine atlas of human PBMC’s could be leveraged to gain a deeper insight into what a patient’s cytokine signature represents at the cellular and gene levels to guide clinicians in prioritizing dysregulated pathways/genes and identifying underlying disease processes in the case of undiagnosed patients.

To do this I present CytoCarto (https://cytocarto.vercel.app), a web application with a simple, interactive user interface which takes as input results of clinical cytokine panel testing and outputs the most likely associated D-SPIN programs, responsible cell types, implicated genes, and diseases/disease mimics (Fig. 4A). CytoCarto (short for Cytokine Cartography) maps the query cytokine profile onto the D-SPIN-generated cytokine map by first taking normalized plasma cytokine measurements and projects them onto both the D-SPIN program network and the GRN (Fig. 4B). Since the 10 million immune cell dataset is comprised of cells from 12 donors of differing age, sex, race, and ethnicity, the program allows the inclusion of this data for the patient and weighs the network responses based on this data. Considering each of the 10 million reference cells is scored for each of the 40 programs, CytoCarto is able to take the program response vector for the patient and assess for immune cell subset enrichment to provide a list of cellular culprits most likely to be responsible for the particular cytokine profile (Fig. 4C). Furthermore, since the subject cytokine profile is propagated over the GRN, predictions on the genes most likely to explain the provided cytokine profile can be identified. This data is displayed as a knowledge-flow network (Fig. 4D) and as simple list (Fig. 4E). Finally, CytoCarto aggregates data from multiple publicly available databases and uses Mitochondrial ROS (mtROS) are recognized to play important roles in immune cell function^31^ and *SOD2* has been found to regulate mtROS-mediated inflammatory responses^32^. Dysregulation of mtROS has been theorized to primary mitochondrial disorders) among the likely disease/mimics. This may be due to limited inclusion of data from this rare patient this to rank possible diseases, disease mimics, pathways, regulatory programs, and cellular contexts which could explain the cytokine profile. In brief, it does this by taking the signed D-SPIN gene scores first generated after propagating the cytokine network across the D-SPIN network, and scoring it against various gene sets from these databases. Statistical corrections are made, disease terms are mapped to MONDO disease identifiers and then ranked. Furthermore, the patient gene vector is propagated through the signed ChEA-KG TF network^30^ to leverage the power of this large GRN to retrieve related gene modules. Finally, human gene signatures from the Perturb-Seqr database were used to identify gene perturbations which phenocopy the patient’s gene response pattern or reverse it. Results of this analysis layer are found in the “Diseases & mimics” tab of CytoCarto (Fig. 4F) and can also be downloaded in full for further analysis (Fig. 4C).

**Figure 4:**
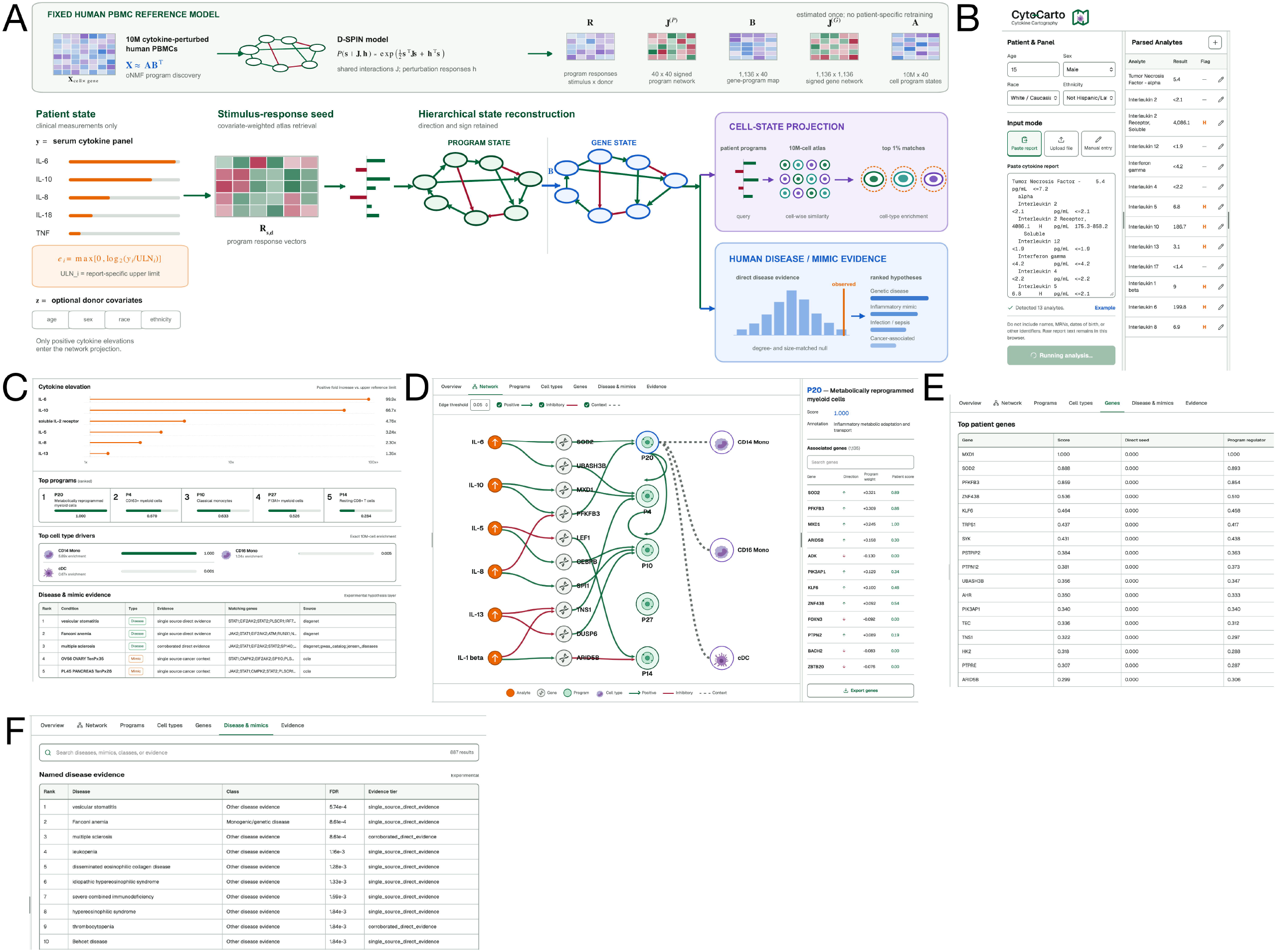
CytoCarto: Data-driven evaluation of human cytokine panels. A. Graphical overview of CytoCarto demonstrating the computational approach used to derive programs, gene, cell type, and disease evidence from a patient’s clinical cytokine panel results. B. Cytokine data input options for CytoCarto. C. Example summary of CytoCarto results showing the top dysregulated cytokines for the patient in the top box, strongest D-SPIN programs linked to this cytokine profile in the second box, most likely responsible immune cell types for the patient’s cytokine profile in the next box, and strongest disease/mimic’s matching the cytokine profile in the final box. The underlying data from the projection of patient’s cytokine profile onto the D-SPIN network can be downloaded with the button located at the top right of the app. D. System network diagram tab demonstrating flow of information from cytokines to genes, to programs, and finally to suspect immune cell types based on input cytokine signature. Edge opacity and width are fixed and both edges and nodes are trimmed to show only the most important. You can click on each program to bring up the right panel showing the genes in that program, how the patient’s cytokine data is predicted to regulate each, and the weight they contributed to the scoring function used to prioritize D-SPIN programs. E. Genes predicted to be dysregulated based on the patient’s cytokine profile. F. Top disease and disease mimics predicted to be related to the patient’s cytokine profile.

To test the utility, accuracy, and power of CytoCarto I input the clinical cytokine profile of a 15-year-old male patient with mitochondrial encephalopathy, lactic acidosis, and stroke-like episodes (MELAS) taken during an episode of undifferentiated inflammation with no identified infectious source and ICU admission for refractory seizures and encephalopathy (Fig. 4B). CytoCarto identified that the most likely program to explain his profile was the “P20-Metabolically reprogrammed myeloid cells” and one of the top genes contributing strongly to his cytokine profile, is *SOD2* (Fig. 4C, E). play a role in mitochondrial sterile inflammation, but a particular role for *SOD2* has yet to be established. It is notable that CytoCarto did not identify MELAS (or any other population in the databases used (Fig. 4F). The fact that CytoCarto correctly identified monocytes as the immune cell type most responsible for sterile inflammation in this patient is supported by recent work in patients with Leigh syndrome (a similar primary mitochondrial disorder) showing the same (C.W. Johnson, presented at the 2026 UMDF meeting). Overall, CytoCarto was able to correctly identify that metabolically deranged monocytes were responsible for this patient’s cytokine profile and further linked an exciting candidate regulatory gene (*SOD2*).

## Discussion

Here, I made use of a recent advance in GRN construction to derive a functionally useful model of human immune cell cytokine responses. In addition to identifying potentially interesting hypothesis, this work provides proof-of-concept for the broader application of custom perturbation screens across immunology. Massive-scale perturbation assays (ie. Perturb-seq) are arriving as the next big step in how we approach complex biological problems. These assays go beyond the exponential effect from adding more parameters to an assay and allow for synergistic scaling of experimental output. When analyzed with evolving network-based computational approaches, such as D-SPIN, they have the potential to reveal important hypothesis-driving findings, such as the importance of *SOD2* in coordinating innate immune cell response to cytokines.

The current limitation to taking such an approach is the complexity and cost of performing large perturbation screens, along with the difficulties inherent to computational analysis and interpretation of such datasets. D-SPIN addresses the later point as demonstrated Jiang et al^4,19^ and here within. The first two points have also recently been addressed through the development of VIPerturb-seq^33^ and the associated GuEST-List genome-wide CRISPRi library. VIPerturb-seq makes use of a prob-based detection method and 10X Genomics FlexV2 platform to drive massive cost and throughput improvements, now making Perturb-seq an assay any lab can run^33^. The last challenge, designing and validating CRISPR libraries is taken care of with the introduction of the GuEST-List library^33^ which makes use of a split barcode design allowing for genome-wide libraries compatible with 10X Genomics FlexV2 chemistry and VIPerturb-seq. I envision the combination of D-SPIN with VIPerturb-seq as being a powerful approach groups will increasingly be turning too in the near future.

There are many future considerations which will need to be addressed when using this combination of technologies. One of which is how granular should cell type annotation be? This is exemplified in this work by a highly metabolically active monocyte subset within the larger CD14 Mono cell type. It is hard to imagine any heuristic threshold that be identified which would apply globally, and the likely answer is that it will greatly depend on your experimental sample and specific question(s). Another important consideration is the perturbation method of choice. D-SPIN’s analysis power is limited by the number of very strong perturbations to the system, and so while the power of D-SPIN scales with the number of perturbations, one must ensure that these perturbations have a potent effect on the system. This also requires one to consider if the system at large needs additional modulation, for example, by varying culture conditions or stimulations to allow a weak perturbation to become powerful.

The introduction of Cytokine Cartography exemplifies the utility of D-SPIN networks and uses this very “basic-science” study of immune cell responses to directly translate into a tool for clinical benefit. In addition to serving as a tool to assist with clinical-grade cytokine panel analysis, CytoCarto could in theory be applied to research-level cytokine panels from any *in vitro* experiments. In this case, results may be even more robust considering the much larger breadth of cytokines which are typically included in multiplex panels. CytoCarto is not without a significant number of limitations discussed below.

The overall study contains a few important limitations. First is the use of fixed, high cytokine doses (which does not take into account that some cytokines have differing effects and cell-type specificity at various concentrations such as IL-2^34^). Additionally, the cytokine stimulations were conducted *in vitro* and there were no combinations of multiple cytokines tested. These limitations all stem from the enormous cost and effort with scaling the number of conditions given the significant cost with an experiment of this nature. Already, this 10 million cell dataset is one of the largest scRNA-seq datasets ever published and is pushing the limits of current analysis technology. CytoCarto is limited in that it includes some heuristic weights in its calculations. It would benefit from training on a large dataset of annotated patient cytokine profiles to refine these weights, and the model at large. CytoCarto is also limited by the reference datasets it compares samples too, as exemplified by a lack of primary mitochondrial disease samples in our test case. As sequencing data on rare disease cohorts continues to expand, CytoCarto will be able to incorporate this data to increase its analytical breadth. Furthermore, CytoCarto is built on data from PBMC’s of a limited number of individuals. This means it cannot account for cytokines and immune responses of tissue resident immune cells, such as macrophages, which are major producers of cytokines and implicated in many inflammatory states. To address this will require performing new functional genomic screens with these difficult to obtain cell types. Finally, CytoCarto is limited by the cytokines available for clinical-grade patient testing given most CLIA-certified cytokine panels remain rather limited in the number of analytes. This can be addressed through the use of large research-grade panels, which if applied to a cohort of patients with undifferentiated autoinflammation, could serve as a powerful hypothesis generating tool to complement paired whole genome/exome sequencing.

## Acknowledgements

I thank Jialong Jiang for assistance with D-SPIN and Rory Tinker for assistance hosting CytoCarto. This work was supported in part through the Minerva computational and data resources and staff expertise provided by Scientific Computing and Data at the Icahn School of Medicine at Mount Sinai and supported by the Clinical and Translational Science Awards (CTSA) grant UL1TR004419 from the National Center for Advancing Translational Sciences.

## Data availability

The 10 million cell scRNA-seq dataset is publicly available through Parse Biosciences. Full output from D-SPIN analysis of this dataset is available here: https://zenodo.org/records/21497681.

CytoCarto is available for use here:

https://cytocarto.vercel.app and its underlying code can be found at: https://github.com/poconnel3/CytoCarto.

