## Supplementary material for "A human cytokine response atlas to reconstruct underlying gene regulatory networks": Supp_figs

### Supplemental Data:

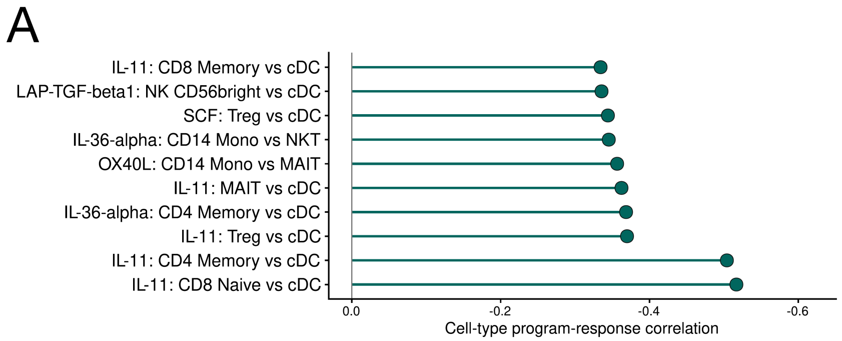


***Supplemental Figure 1: Negative correlations of cytokine responses between cell types.***

*A. Lollipop plot of the most negative correlations between cell types for each cytokine.*

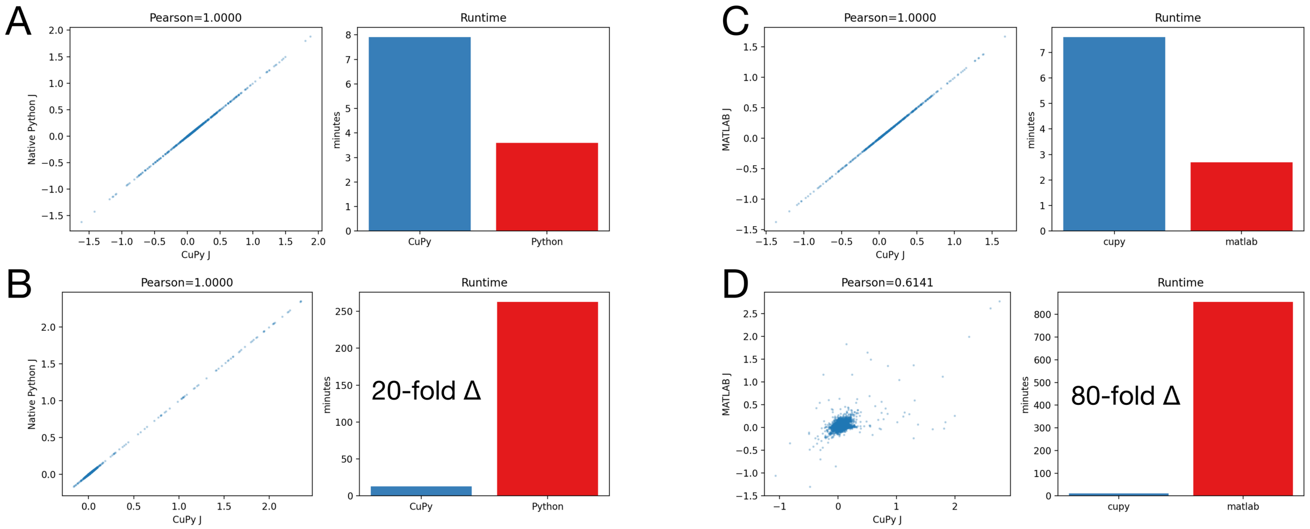


***Supplemental Figure 2: Validation of GPU optimization of D-SPIN.***

*A. Left, correlation of J values between standard python implementation of D-SPIN and GPU-D-SPIN (AKA CuPy) on program-level network calculations using a 500,000 cell subset of the cytokine atlas dataset. Right, bar plot of run times. B. Same as (A), but showing results from the gene-level GRN D-SPIN calculation. C. Left, correlation of J values between MATLAB implementation of D-SPIN and GPU-D-SPIN on program-level network calculations using a 2 million cell subset of the cytokine atlas dataset. Right, bar plot of run times. D. Same as (C), but showing results from the gene-level GRN D-SPIN calculation.*
